# Pik3ip1/TrIP Regulation of PI3K Restricts CD8 T Cell Anti-Tumor Immunity

**DOI:** 10.1101/2025.07.10.664200

**Authors:** Benjamin M. Murter, Lawrence P. Kane

## Abstract

**Background:** The protein PI3K-interacting protein (PIK3IP1), or transmembrane inhibitor of PI3K (TrIP), is highly expressed by T cells and can modulate PI3K activity in these cells. Several studies have also revealed that TrIP is rapidly downregulated following T cell activation and can play important roles in T cell differentiation.

**Methods:** We generated mice with CD8-specific TrIP deficiency. We then implanted these mice, and Cre-only control animals, with B16 melanoma or MC38 colon carcinoma tumors. Tumor growth and anti-tumor immunity were then followed. We also assessed the effects of TrIP deficiency on transcriptional programs in CD8 T cells stimulated in vivo or derived from tumor-bearing mice.

**Results:** We found that activated TrIP KO CD8 T cells display an increased inflammatory transcriptional profile in the absence of TrIP. Consistent with these effects, we also found that knockout of TrIP specifically in CD8 T cells resulted in reduced growth of syngeneic tumors. When characterizing the tumor-infiltrating cells, TrIP KO led to an increase in the number of tumor-infiltrating T cells, as well as a delay in the acquisition of an exhausted phenotype, based on phenotypic and transcriptomic analyses. Finally, our data suggest that TrIP regulates the diversity of T cell clonal responses to tumors, since we observed an increase in the number of distinct T cell clonotypes responding to a tumor neoantigen.

**Conclusions:** Taken together, our findings demonstrate that TrIP intrinsically restricts the CD8 T cell response to tumors, and that targeting TrIP may augment the anti-tumor response in a way that is distinct from established checkpoint therapies.

## Background

CD8 T cells are critical for the adaptive immune response to viral infection and tumors. In the absence of the clearance of antigen, as often observed in chronic viral infection or tumors, persistent stimulation results in cytotoxic CD8 T cells acquiring an “exhausted” phenotype. This state is characterized by increased expression of inhibitory (aka “checkpoint”) receptors (e.g. PD-1, CTLA-4, Tim-3, Lag-3), as well as reduced effector cytokine production (e.g, IFNγ, IL-2) and proliferation [1–3]. Importantly, exhausted T cells are not completely dysfunctional, as they still retain many of their effector functions such as cytotoxicity and some cytokine production, albeit at significantly lower levels. Advances in cancer immunotherapy have revealed that blockade of receptors like PD-1/PD-L1 can reinvigorate a proportion of the exhausted CD8 T cells and expand their cytotoxicity and cytokine production [4]. Understanding why only a subset of these ‘exhausted’ cells respond to checkpoint blockade is an area of intense investigation. One model posits that the differentiation fate of CD8 T cells, at least within the context of tumor antigen, is imprinted early upon antigenic stimulation, via distinct epigenetic signatures [5]. T cell help during initial antigen encounter in the lymph nodes is increasingly believed to alter the epigenetic landscape of CD8 T cells, promoting certain gene expression profiles and favoring or repressing individual lineage commitments [6]. Current immunotherapy modalities are limited by the dysfunctional state of most T cells in the TIL. Therefore, an increasing focus of the field is to understand how to improve or promote an activated T cell program that results in a more effective, less exhausted, anti-tumor T cell tumor response.

Activation and acquisition of cytotoxic effector activity by CD8 T cells require multiple signals delivered through the TCR/CD3 complex (Signal 1), co-stimulatory molecules like CD28 (Signal 2) and cytokine receptors (Signal 3) [7]. One of the biochemical pathways that has been linked to all three of these signals is Class I phosphatidylinositol 3-kinase (PI3K) [8, 9]. The Class I PI3Ks comprise heterodimeric regulatory and catalytic subunits, of which there are multiple isoforms (α, β, γ, δ). PI3Ks generate the lipid second messenger PIP_3_, which directly or indirectly aids in the activation of multiple downstream proteins, including the kinases Itk, PDK- 1, Akt, mTOR and S6K. As a positive factor in cellular activation and survival, this pathway must be carefully controlled; indeed, alterations in PI3K can lead to proliferative dysregulation and even cancer. Most known negative regulators of PI3K, including PTEN and SHIP, act by directly dephosphorylating PIP_3_ [10].

Recent studies described a membrane-bound inhibitor of PI3K, *PIK3IP1* or TrIP (transmembrane inhibitor of PI3K), which operates upstream of any previously described negative regulators of PI3K [11]. TrIP was initially discovered as a novel kringle domain-containing protein, the cytoplasmic domain of TrIP (PIK3IP1) was found to have significant homology to the inter-SH2 domain of the p85 PI3K adaptor proteins [11, 12]. This work suggested TrIP imparts its regulation through directly interacting/inhibiting with the PI3K p110/p85 heterodimer. More recent reports have shown that the expression of TrIP is particularly high on naïve/resting CD8 T cells, and that it is downregulated in a manner proportional to TCR signal strength [13]. A role for TrIP in T cell differentiation was demonstrated through *in vitro* helper T cell skewing experiments, in which the absence of TrIP promoted a generally more inflammatory Th1 phenotype and impaired the induction of peripheral Treg [14]. These data were supported by an *in vivo* mouse model of Listeria monocytogenes infection, where mice with T cell-specific knockout of TrIP (using CD4-Cre) showed a significantly lower bacterial burden compared to wild-type mice.

Here we have examined how TrIP regulates early activation in CD8 T cells and the impact of TrIP on T cell activation and subsequent anti-tumor immune response. While a recent study has shown germline TrIP KO mice displayed anti-tumor resistance, although the mechanisms of these effects are largely unclear [15]. In this study we detail how the absence of TrIP regulation on naïve CD8 T cells alters their gene expression pattern, resulting in an increased inflammatory phenotype. Furthermore, we found that this increased inflammatory phenotype is sufficient to lower tumor burden in a CD8-specific knockout of TrIP. We investigated the tumor immune infiltrate and found the absence of TrIP expression led to greater T cell infiltration, driven by the CD8 T cell compartment. Mechanistically, we present data suggesting that the absence of TrIP regulation during early activation events within the tumor draining lymph nodes (TDLNs) alters the clonal diversity of the CD8 T cell response.

## Methods

### Mice

E8i-cre (Strain #008766), CD4-cre (Strain #:022071), and P14 TCR transgenic (Strain #037394) mice were originally obtained from The Jackson Laboratory and bred at the University of Pittsburgh. The TrIP conditional knockout mouse strain were generated by inGenious Targeting Laboratory Inc., using C57BL/6 ES cells. LoxP sites were inserted into the TrIP gene (*Pik3ip1^fl/fl^*), targeting the removal of exons 2-5 via Cre-mediated recombination. For peptide stimulation studies, we employed the P14 TCR transgenic strain that carries a rearranged T cell receptor specific for the gp33 peptide from LCMV (Jackson; Strain #037394). All strains were backcrossed at least nine generations to C57BL/6J (Jackson; Strain #000664). All mice were bred in-house under SPF conditions, and experimental mice were either littermates or housed in the same facility and same room. All experiments were conducted on mice 6-8 weeks of age, with age and sex-matched groups. All animal procedures were conducted in accordance with NIH and University of Pittsburgh IACUC guidelines.

### Antibodies and Staining

For extracellular staining, single cell suspensions were stained at 4°C for 30 minutes with Live/Dead + antibody cocktail containing anti-CD16/CD32 (Fc Shield; Tonbo Biosciences: clone 2.4G2) resuspended in PBS. For intracellular staining, cells were fixed/permeabilized using the eBioscience Foxp3/transcription factor staining buffer set (00-5523-00), as per manufacturer instructions. Following fixation, intracellular staining antibodies were resuspended in 1x permeabilization buffer and used to stain at 4°C for 30 minutes.

The following antibodies and dyes were used: anti-CD90.2 (BD Biosciences; clone 53- 2.1), anti-CD8α (BD Biosciences; clone 53-6.7), anti-CD44 (BD Biosciences; clone IM7), anti-TCRβ (BD Biosciences; clone H57-597), anti-CD4 (BioLegend; clone RM4-5), anti-CD62L (BioLegend; clone MEL-14), anti-CD25 (BD Biosciences: clone 7D4), anti-Foxp3 (Invitrogen; FJK-16s), anti-pS6(235/236) (Cell Signaling; clone D57.2.2E), anti-CD69 (BioLegend: clone H1.2F3),Live/dead staining was performed using the Zombie NIR fixable viability kit (BioLegend; 423105). All flow cytometry was performed on a 5-laser Cytek Aurora and flow cytometry data were analyzed with FlowJo (v10.10.0).

### Reagents and Media

TDLNs harvested from mice were mechanically disrupted and filtered to single cell suspensions. Red blood cells were removed via 1x RBC lysis buffer (eBioscience,00-4333-57), according to manufacturer instructions. Tumor (TIL) and TDLN re-stimulations were done in complete RPMI (cRPMI) media (cRPMI supplemented with 10% BGS, 1% penicillin/streptomycin, 1% L-glutamine, 0.05 mM 2-mercaptoethanol, 1x Non-Essential Amino Acids, 1% HEPES, and 1% sodium pyruvate). Anti-CD3 (clone:145-2C11; Tonbo Biosciences 50-201-4837); and anti- mCD28, clone 37.51; BioLegend: 102101) were used for plate-coating at the concentrations indicated in the figure legend.

### Tumor Models

Syngeneic MC38 colon carcinoma cell line was obtained from Dr. Dario Vignali. Tumor cells lines were maintained in (cDMEM) DMEM containing 10% Bovine Growth Serum (BGS), 1% penicillin/streptomycin, and 1% L-glutamine. For tumor challenge experiments, 5x10^5^ MC38 cells were injected subcutaneously (SubQ) into the mouse flank, and tumor growth was followed for 2-four weeks. Tumors were measured every 2-3 days via digital calipers, recording the length and width with perpendicular measurements. Tumor size reported as ‘Tumor Area (mm^2^)’ multiplying the two measurements together. For tissue harvests, excised tumors were roughly chopped and then digested using collagenase D and 0.2 mg/ml DNase I, type IV for 40 min at 37°C with constant gentle shaking in Miltneyi C-tubes, using a GentleMACS tissue dissociator (Miltenyi). Reaction was then quenched with 5x cold cDMEM, filtered through 70 μM cell strainer, and washed. RBC lysis is performed with a subsequent wash. The resulting single-cell suspension was counted and then used for the downstream flow cytometry staining and analysis.

### Tetramers

Rpl-18* tetramers, an H2-Kb class I-presented neoepitope from the 60S ribosomal protein L18 [16] (peptide sequence: KILTFDRL), were obtained from NIH tetramer Core. Staining performed with other surface staining.

### RNA sequencing

For the *in vitro* stimulation experiment: Splenocytes were harvested from *Pik3ip1^fl/fl^* CD4^cre^ P14^tg^ (KO) and CD4^cre^ P14^tg^ (WT) mice. All conditions received rIL-2 (50U/ml) and rIL-12 (2ng/ml) to promote an effector differentiation program. Cells were cultured under three different conditions; no stimulation, stimulated, and stimulated in the presence of the PI3Kδ inhibitor (IC87114, 10uM). Stimulation was performed using 100 ng/ml of cognate WT gp33 peptide. following 24hrs in the indicated culture conditions, activated (CD44^+^) CD8 T cells were sorted from each population and performed bulk RNAseq. For the day 10 Rpl18 RNAseq experiment: Tissues were harvested, processed, and stained as detailed above. FACS-sorting was performed using a BD FACS-Aria.

Full-length cDNA was generated with the Takara SMART-Seq mRNA HT kit (Takara: 634796) according to the manufacturer’s instructions, with an additional 0.5 ml TCR specific oligo-dT primer from the Takara SMARTer Mouse TCR kit (Takara: 634403). Between 425-1000 cells were sorted into 12.5 µl of lysis buffer and snap frozen on dry ice prior to storage at -80°C. Immediately prior to cDNA generation, samples were incubated at 72°C for 3 min then held on ice. cDNA was generated at 42°C for 90 minutes, followed by 15 cycles of SMART whole genome amplification. cDNA was assessed for quality using an Agilent Fragment Analyzer 5300. Whole transcriptome libraries were generated with the Illumina DNA Prep kit (Illumina: 20060059) according to the manufacturer’s instructions, using 5 ng of input cDNA for each sample and 11 cycles of indexing PCR were completed using Illumina DNA/RNA UD Indexes. TCR-seq libraries were generated using 3 ng of input cDNA and 10 cycles of PCR1 amplification using TCR A and B specific primers, and 20 cycles of indexing PCR2 using indexed TCR specific primers (Takara: 634403). Library quantification and assessment was done using a Qubit FLEX fluorometer and an Agilent TapeStation 4150. Libraries were normalized and pooled to 2 nM by calculating the concentration based off the fragment size (base pairs) and the concentration (ng/ml) of the libraries. Whole transcriptome sequencing was performed on an Illumina NextSeq 2000, using a P3 200 flow cell with read lengths of 2x101bp, with a target of 40 million reads per sample. TCR sequencing was performed on an Illumina NextSeq 2000, using a P1 600 flow cell with read lengths of 2x301bp, with a target of 10 million reads per sample.

Sequencing data was demultiplexed by the on-board Illumina DRAGEN FASTQ Generation software. Data analysis was performed using CLC Genomics Workbench, Partek Flow, and/or Ingenuity Pathway Analysis Suite software, differential gene analysis by DESeq2. TRUST4 was used for the TCRseq repertoire analysis. Datasets are available from the Gene Expression Omnibus (GEO), accession numbers: GSE302184, GSE302185 and GSE302196.

## Results

### RNAseq reveals an enhanced inflammatory phenotype of TrIP KO CD8 T cells

We recently reported that TrIP is highly expressed on naïve/resting CD8 T cells and that its ecto domain is proteolytically cleaved from the cell surface after activation through the TCR [13]. Additionally, previous work from our lab and others revealed that TrIP-deficient CD4 helper T cells are skewed towards an increased inflammatory phenotype under polarizing conditions *in vitro* [14]. Thus, we wanted to investigate how the absence of TrIP expression alters the gene expression and function of CD8 T cells after stimulation. Its unique position upstream of PI3K suggests that TrIP deletion may result in a transcriptional profile distinct from that of other known regulators of this pathway (e.g. PTEN). Additionally, shifts in metabolic products and intermediates (i.e. acetyl-CoA) are known to modify the gene expression programs of T cells [17], and TrIP has already been reported to influence the metabolic profile of T cells [18]. We bred conditional TrIP knockout mice (*Pik3ip1^fl/fl^*) to mice expressing Cre in CD8 T cells (E8i-Cre) (Jackson; Strain #008766)and P14 TCR transgenic (Tg) mice expressing a receptor specific for the H-2D^b^ gp33-41 epitope from LCMV (Jackson; Strain #037394). Since TrIP expression is high on naïve cells, we evaluated the transcriptome before and after 24h of activation with the gp33-41 peptide, in the presence or absence of the selective PI3K p110δ inhibitor IC87114 (PI3Ki) [19]. The latter was included to determine what extent the effects of TrIP deletion are dependent on PI3K.

As shown in **figure 1**, the absence of TrIP expression led to significant differences in the expression of inflammatory gene expression across three conditions: naïve, stimulated and stimulated with PI3K inhibitor. Principal component analysis (PCA) revealed a large separation across PC1 between samples, indicative of the presence or absence of TCR stimulation in these samples (**figure 1A**). TrIP deficiency resulted in major transcriptional changes, as the PCA plot shows that each TrIP KO group clustered separately from their WT counterparts across PC2 (**figure 1A**). To understand the genes driving these differences, we looked for differentially expressed genes (DE genes: absolute fold change ≥2, and p-value <0.05) across each condition. The TCR stimulation-alone condition resulted in the highest total number of DE genes pik3ip1 (1858), the majority of which were upregulated (1358 genes upregulated, 498 genes downregulated). The unstimulated and the TCR stimulation + PI3K inhibitor conditions each resulted in over 800 DE genes. These data demonstrate that TrIP KO CD8 T cells are transcriptionally different from their WT counterparts, regardless of pMHC-TCR stimulation. Intriguingly, pharmacologic inhibition of the PI3K pathway led to a significant reversal of the effects of TrIP KO (gold and green symbols) (**figure 1A**). To better understand which pathways are enriched in the absence of TrIP expression, we performed gene set enrichment analysis (GSEA) on data from the stimulated cells (**figure 1B**). Interferon response gene sets showed the greatest enrichment, but cytokine related pathways such as IL-2/Stat5 signaling were also enriched. This is interesting, as IL-2 is known to stimulate the PI3K pathway [20], suggesting there may be differential cytokine responses in the absence of TrIP. Although there were significant differences in many pathways and genes, not all DE genes were shared across the treatment groups. A heat map with relative expression of selected genes is shown in **figure 1C**. Quantitative expression of several effector-associated genes is also shown, in **figure 1D**. Since the transcription factor FOXO1 is a known target of PI3K/Akt signaling in T cells [21, 22], we also performed targeted gene set enrichment analysis, using a curated set of genes activated or repressed by FOXO1 [23].

**Figure 1:**
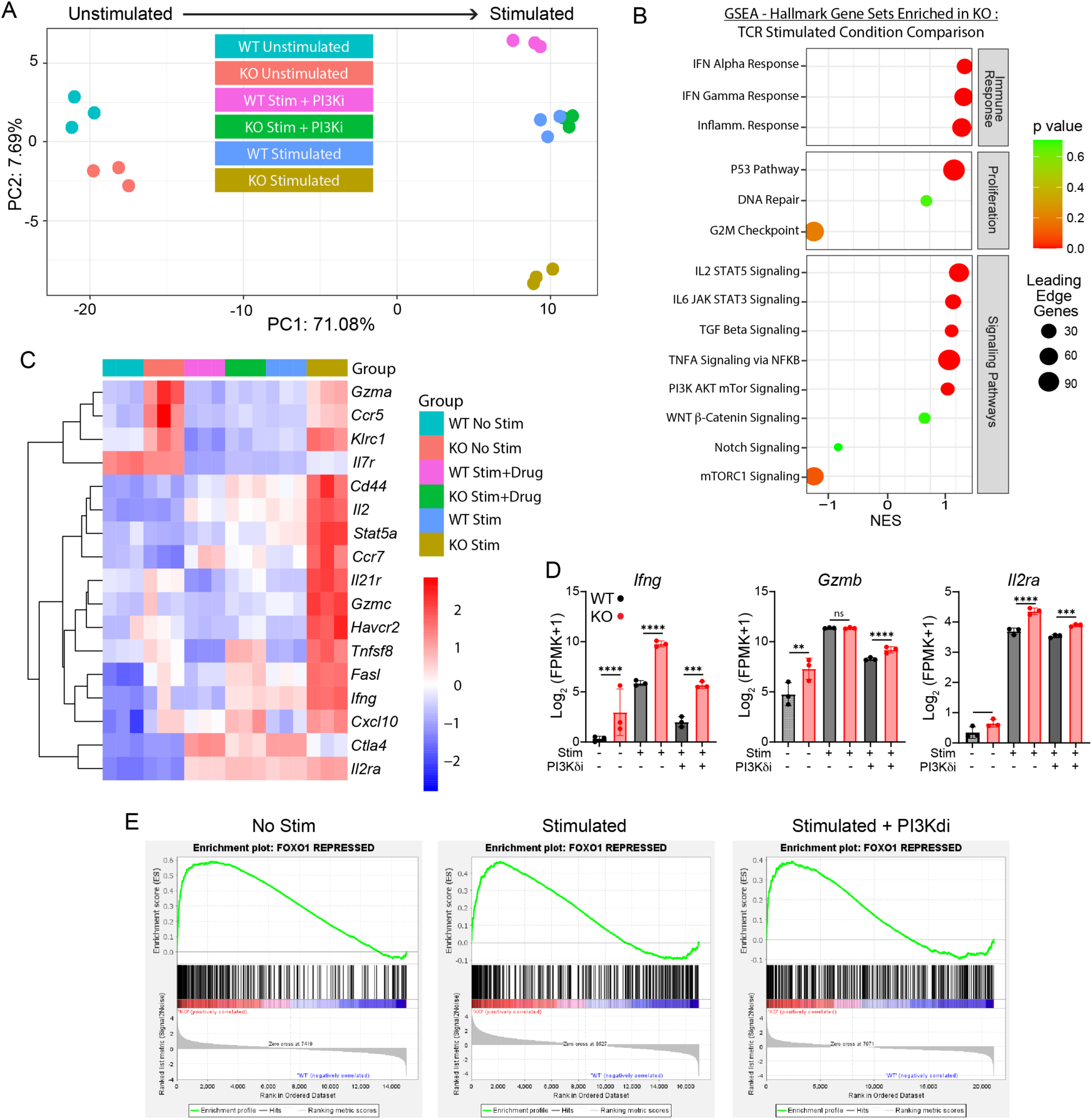
Absence of TrIP promotes an enhanced inflammatory transcriptional profile in activated CD8 T cells. Splenocytes from WT and CD8-specific TrIP knockout P14 mice were cultured for 24 hrs under 3 different conditions; Unstimulated (50 U/ml rIL-2 + 2 ng/ml rIL-12), Stimulated (100 ng/ml HiAff gp33 peptide + 50 U/ml rIL-2 + 2 ng/ml rIL-12), and stimulated + PI3Ki (10 μM IC87114 + 100 ng/ml high affinity gp33 peptide + 50 U/ml rIL-2 + 2 ng/ml rIL-12). After 24 hr stim., CD8 T cells were sorted out and bulk RNAseq was performed. (**A**) Principal component analysis (PCA). (**B**) Gene set enrichment analysis (GSEA) of the stimulation condition, showing gene sets enriched in the TrIP knockout. (**C**) Heatmap of normalized counts of selected genes. (**D**) Bar graphs of normalized gene counts across all conditions for the selected inflammatory genes: *Ifng*, *Gzmb*, *Il2ra*. (**E**) Gene set enrichment analysis of differentially expressed genes vs. a FOXO1-repressed gene set. Ordinary one-way ANOVA was applied for statistical comparisons (p: *<0.05, **<0.01, ***<0.001, ****<0.0001).

### CD8-specific TrIP knockout leads to lower syngeneic tumor burden

To date, only one study has investigated the role of TrIP regulation in the context of tumors where it was demonstrated that germline TrIP KO mice displayed enhanced anti-tumor immunity [15]. However, the direct mechanisms behind these effects are still not clear. Based on this finding and the above transcriptomic data, we wanted to investigate how the absence of TrIP expression specifically on CD8 T cells alters the anti-tumor immune response. For these studies, we used the conditional knockout mouse described above (without the TCR Tg). We performed syngeneic tumor challenges with either immunotherapy-sensitive (MC38) or immunotherapy-resistant (B16-F10) tumors, which were implanted subcutaneously on the rear flank (**figure 2A**). When WT (Cre-only) and CD8-specific TrIP KO mice were challenged with the MC38 tumor line, we observed that knockout of TrIP expression in CD8 T cells alone was able to significantly reduce the tumor burden compared to WT controls (**figure 2B**). Even more impressively, the TrIP knockout mice also showed resistance to the more aggressive, and less immunogenic, B16F10 tumors, compared to Cre-only controls (**figure 2C**).

**Figure 2:**
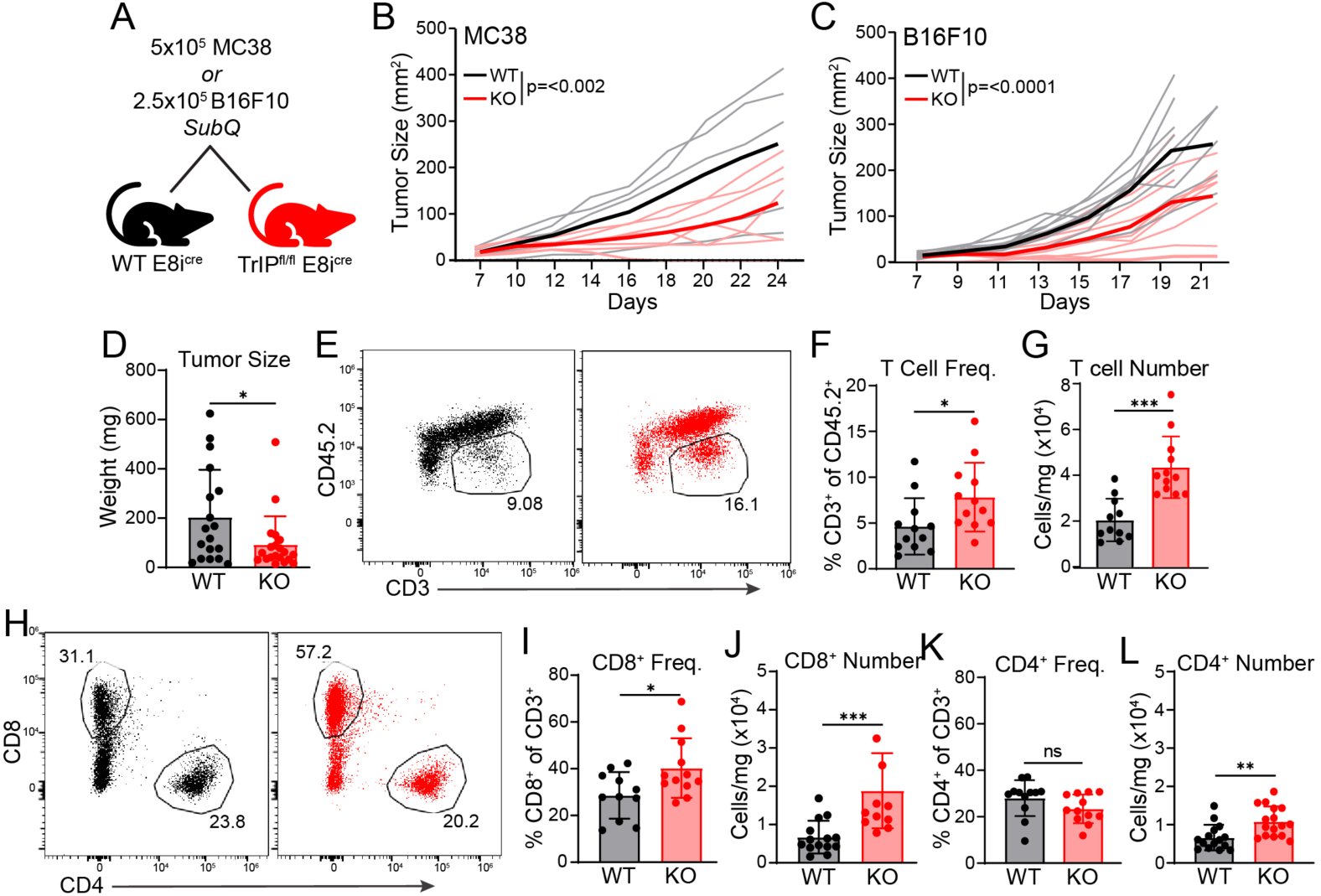
Increased frequency of intratumoral TrIP KO CD8 T cells at day 14. MC38 or B16-F10 cells were injected subcutaneously on the rear flank of *Pik3ip1^fl/fl^* x E8I^Cre^ (KO) or E8I^Cre^ (WT) mice; tumors were measured via digital calipers every 2-3 days. Tumor size expressed as mm^2^ (calculated from LxW). (**A**) Experimental design. (**B-C**) Tumor growth curves after injection of MC38 (B) or B16-F10 (C) tumors. Ordinary two-way ANOVA was applied for statistical comparisons (p: *<0.05, **<0.01, ***<0.001, ****<0.0001). Tissues were harvested on day 14 and immune infiltrate was analyzed by flow cytometry. **(D**) Tumor size, by weight in milligrams (mg). (**E**) Flow plots of CD3 T cell gate of both WT (black) and knockout (red). (**F**) Frequency of CD3 T cells of the total CD45.2^+^ within the tumor. (**G**) Extrapolated cell counts of CD3 T cells, normalized to tumor weight in mg. (**H**) Flow plots of CD4 and CD8 T cell gates of both WT (black) and knockout (red). (**I**) Frequency of CD8 T cells of the total (CD3^+^) T cells within the tumor. (**J**) Total counts of CD8 T cells, normalized to tumor weight in mg. (**K**) Frequency of CD4 T cells of the total CD3^+^ within the tumor. (**L**) Extrapolated cell counts of CD4 T cells, normalized to tumor weight in mg. Students T-test statistical analysis applied for statistical comparison (p: *<0.05, **<0.01, ***<0.001, ****<0.0001).

### Phenotype of WT and TrIP KO CD8 T cells on day 14 after tumor challenge

We next sought to understand how the absence of TrIP on CD8 T cells alters the tumor immune response. Thus, tumors were harvested fourteen days after implantation and the tumor-infiltrating lymphocytes were analyzed by flow cytometry. Consistent with data discussed above the TrIP conditional knockout mice had smaller tumors (by weight) (**figure 2D**), but no differences in the size (cell count) of the tumor draining lymph nodes (TDLN). Analysis of the TIL revealed a higher frequency of CD3 T cells in the TrIP knockout tumors (**figure 2E,F**). This difference was driven by an increase in the CD8 T cell compartment (**figure 2H,I,J**), with a corresponding reduction in the proportion of CD4 T cells (**figure 2H,K**). In addition to T cell frequency, normalizing these cell counts to tumor weight revealed TrIP knockout TIL to contain significantly more T cells (CD3^+^) overall (**figure 2G**). While the total number of CD4 T cells was increased (**Fig. 2K**), CD8 T cells showed the greatest increase, compared to WT T cells (**figure 2I**).

With the removal of a negative regulator of T cell activation like TrIP, there were concerns that these cells may have an accelerated progression towards the acquisition of an ‘exhausted’ phenotype. In fact, we observed no differences in the expression of PD-1 and Tim-3 on the CD8 T cells within the TIL (**figure 3A,B**). Despite similar PD-1/Tim-3 expression, the TrIP knockout CD8 T cells had significantly lower expression of the transcription factor TCF-1 (**figure 3C,D**). TCF-1 expression is associated with a more stem-like T cell state, suggesting that TrIP knockout T cells acquired a more terminally differentiated effector state within the TIL without grossly altering their exhaustion profile, at least at this timepoint. Broader investigation of other activation and proliferation markers (i.e. Ki-67, KLRG1, Slamf6, and others) also failed to reveal any major differences between the KO and WT T cells within the TIL.

**Figure 3:**
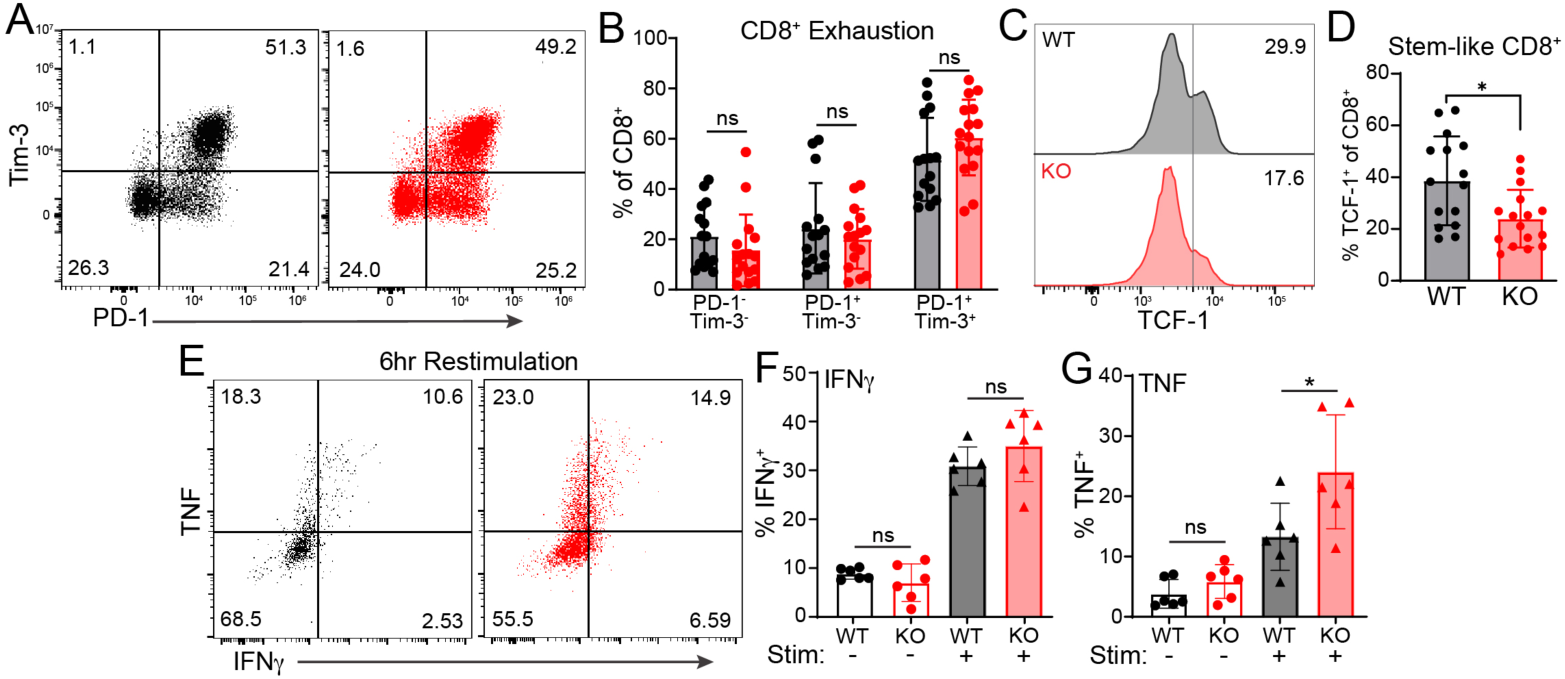
CD8 T cells from day-14 MC38 tumors from *Pik3ip1^fl/fl^* E8i^cre^ (KO) and E8i^cre^ (WT) mice. (**A**) Flow plots of the PD-1 vs Tim-3 expression on the CD8 T cells from the TIL of both WT (black) and knockout (red). (**B**) Quantification of PD-1 vs Tim-3 expression on TIL CD8 T cells. (**C**) Histograms of TCF-1 expression on CD8 T cells within the TIL. (**D**) Quantification of TCF-1^+^ CD8 T cells within the TIL. For cytokine analysis, cells were restimulated with 2 μg/ml plate-coated anti-CD3 for 6 hrs in the presence of Brefeldin A (BFA) and monensin, analyzed via flow cytometry. (**E**) Flow plots of TNF and IFNγ expression following 6 hrs of stim. in both WT (black) and KO (red). (**F**) Quantified total IFNγ expression direct ex-vivo (no stim) and after 6 hrs of TCR stim. (**G**) Quantified total TNF expression ex- vivo (no stim) and after 6 hrs of TCR stim. Students T-test statistical analysis applied for statistical comparison (p: *<0.05, **<0.01, ***<0.001, ****<0.0001).

Although no differences were seen in the expression of exhaustion markers on TrIP KO CD8 T cells, exhausted cells are known to retain some cytokine production and effector activity. This, in addition to the differential expression of TCF-1, led us to investigate if there are any functional differences between the KO and WT CD8 T cells from the TIL. While cytokine expression directly *ex vivo* was low, upon TCR restimulation both WT and KO show increased TNF and IFNγ, with the knockouts showing significantly higher TNF compared to WT (**figure 3E-G**). With these observations, we were still left with an incomplete understanding of the mechanisms of how the absence of TrIP could lead to the increased T cell phenotype we observed within the TIL at D14. With the high activation (% of CD44^+^) of the Rpl18^+^ cells we observed in these D14 TDLN’s, it is possible that we need to look earlier after tumor challenge to see the early activation events where TrIP regulation may have the greatest influence.

### TrIP knockout T cells are less exhausted on day 10 after tumor implantation

Building on the day 14 data, we investigated the response four days earlier, i.e. on day 10 post-implantation, in both the tumor and the TDLN, using the previously discussed Rpl18 tetramer to track antigen specific responses. As described above, we implanted TrIP^CD8^ KO and WT mice with MC38 tumors, but in this case harvested the tumors and TDLN ten days later (**figure 4A**). The tumor sizes were much smaller overall at this earlier timepoint, and there were no significant differences between KO and WT mice (**figure 4B**). Similarly, we also observed no difference in the TDLN size, based on cell count (**figure 4C**). One of the prominent phenotypes we found within the TrIP^CD8^ KO TIL on day 14 was an increased frequency and total number of T cells, driven by the CD8 T cell compartment. This increased T cell frequency appears to have not yet established by day 10, as we saw no differences in the total number of T cells (CD90.2^+^) or in the frequencies of CD8 or CD4 TIL T cells (**figure 4D-H**). We wondered if the absence of TrIP would alter the antigen-specific response in these tumors. However, Rpl18-specific CD8 T cells were detected in the TIL of both KO and WT tumors at comparable frequencies (**figure 4I- J**). Although we found no significant differences in the frequency of CD8 T cells in the TIL, TrIP knockout T cells did have lower exhaustion marker expression (**figure 4K-L**). The greatest difference was seen in the PD-1^+^Tim-3^+^ double positives, suggesting the absence of TrIP expression may slow the progression towards exhaustion, at least initially as we know TrIP KO CD8 T cells expressed similar levels of those markers by day 14. In fact, the WT and KO CD8 shared similarly low levels of TCF-1 expression even at this early timepoint (**figure 4M-N**). Despite higher expression of exhaustion markers, WT CD8 T cells displayed a greater frequency of Ki-67 expression, a marker of proliferation (**figure 4O-P**). We also investigated the cytokine production capacity of these CD8 T cells isolated from day-10 TIL upon restimulation. Following six hours of restimulation, there was robust TNF and IFNγ production from both WT and KO cells (**figure 4Q**). Unlike on day 14, TrIP knockout CD8 did not display an increase in cytokine production compared to the WT by either frequency (% positive) or MFI at day 10 (**figure 4Q**).

**Figure 4:**
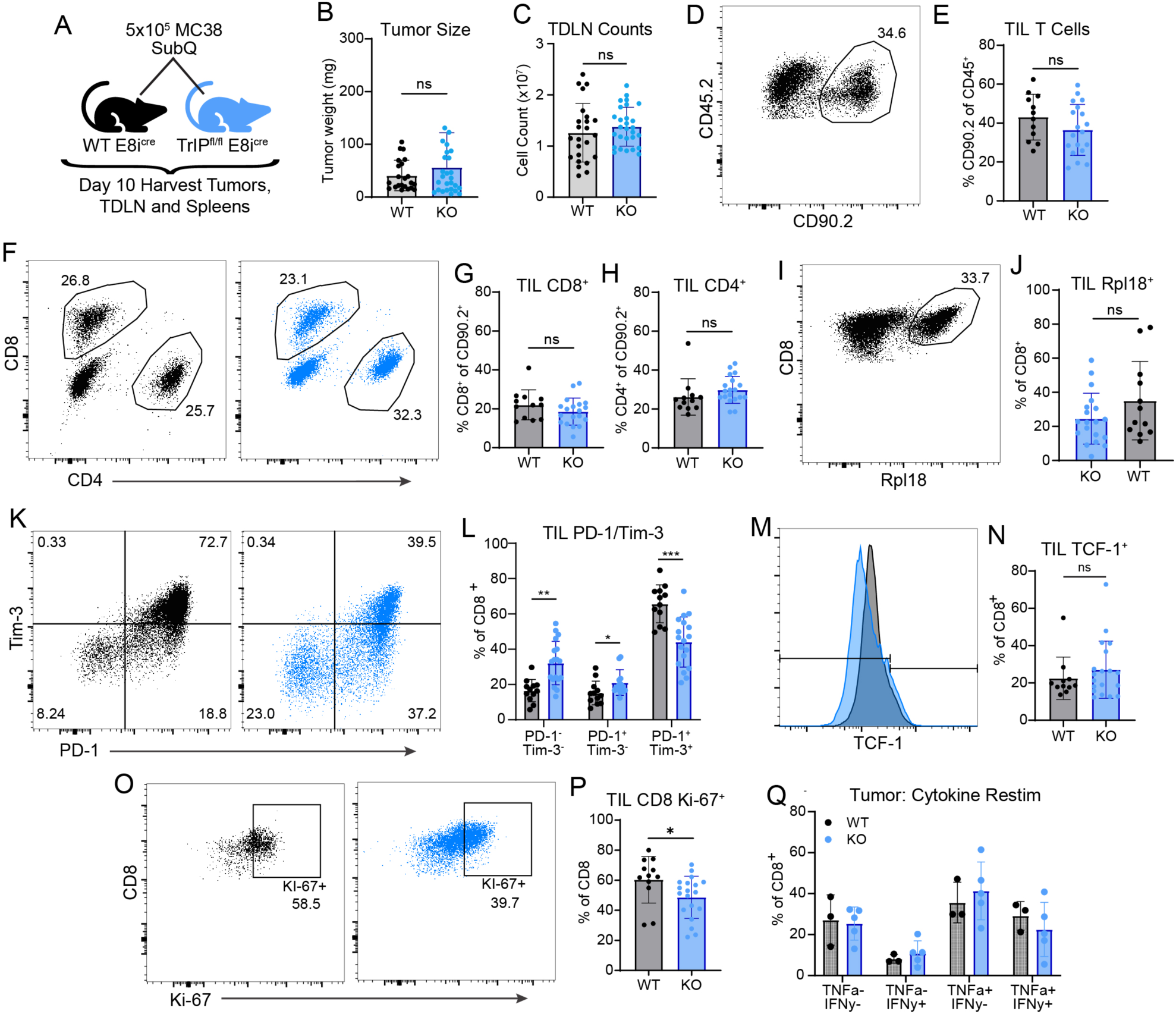
TrIP knockout CD8 T cells are phenotypically less exhausted within the TIL of day-10 MC38 tumors. (**A**) Experimental design. MC38 cells were injected subcutaneously in the flank of *Pik3ip1^fl/fl^* E8i^cre^ (KO) and E8i^cre^ (WT) mice. Tissues were harvested 10 days later and immune infiltrates were analyzed by flow cytometry. (**B**) Tumor size, by weight in milligrams (mg). (**C**) Total lymphocyte counts of the tumor-draining lymph nodes. (**D-E**) Flow plots (D) and quantitation (E) of CD90.2^+^ T cells of the total CD45.2^+^ cells within the tumor. (**F-H**) Flow plots and quantitation of CD4 and CD8 T cells from WT (black) and TrIP^CD8^ knockout (blue), gated on total (CD90.2^+^) T cells within the tumor. (**I-J**) Representative flow plot (I) and quantitation (J) of Rpl18^+^ tetramer staining vs. CD8 cells within the TIL. (**K-L**) Flow plots (K) and quantation (L) of PD-1 and Tim-3 expression on the CD8 T cells of WT (black) and TrIP knockout (blue) CD8 T cells. (**M**-N) Histogram of TCF-1 expression on total CD8 T cells within the TIL. (**N**) Frequency of TCF-1^+^ CD8^+^ T cells within the TIL. (**O-P**) Flow plots (O) and quantitation (P) of Ki-67 expression in CD8 T cells of WT (black) and TrIP knockout CD8 T cells (blue). (**Q**) CD8 T cells from TIL were stimulated with plate-coated anti-CD3 (2 μg/ml) for 6hrs in the presence of brefeldin A (BFA) and monensin, then stained for TNF and IFNγ. Students T-test statistical analysis applied for statistical comparison (p: *<0.05, **<0.01, ***<0.001, ****<0.0001).

We also used the Rpl18 tetramer to investigate the antigen specific response at the earlier day 10 timepoint in the tumor-draining lymph node. Similar to day 14, the day 10 TDLNs of WT and KO mice were similar in size (**figure 5A**). The total T cell (CD90.2^+^) frequency was also similar between the groups (**figure 5B**). We did observe a small (∼5%), but statistically robust, shift in the proportion of CD8 vs. CD4 T cells, with TrIP knockout T cells having a higher frequency of CD8 T cells, compared to WT (**figure 5C-E**). Although there was no difference in Rpl18*-specific frequency in the tumor at day 10, we saw a significant reduction in the KO tetramer^+^ cells in the TDLN (**figure 5F-G**). This shift may be related to the overall increase in CD8 T cell frequency, suggesting a difference in the breadth of responding T cell clones between the KO and WT.

**Figure 5:**
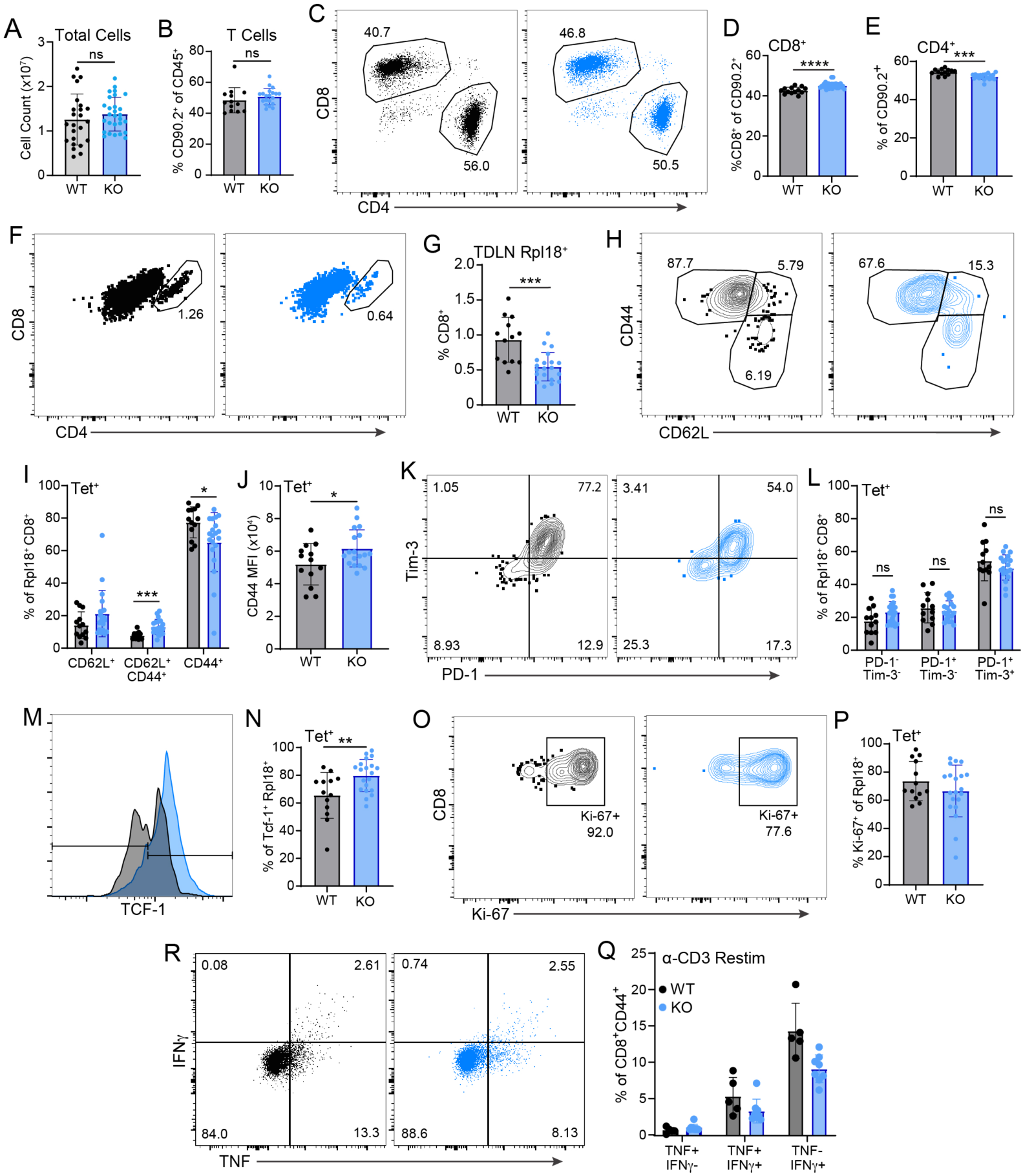
Effects of CD8-specific TrIP KO on TDLN T cells at day 10 after tumor implantation. Tumor draining lymph nodes (TDLNs) were harvested from *Pik3ip1^fl/fl^* E8i^cre^ (KO) and E8i^cre^ (WT) mice on day 10 following MC38 tumor injection. (**A**) Total cell counts. (**B**) Frequency of CD90.2^+^ T cells in the TDLN. (**C-E**) Flow plots (C) and quantitation (D-E) of WT (black) and TrIP KO (blue) CD4 and CD8 T cells. (**F**) Flow plots of WT (black) and KO (blue) CD4 vs. CD8 T cells, based on the Rpl18 tetramer+ gate. (**G**) Frequency of Rpl18 tetramer+ CD8 T cells. (**H-I**) Flow plots (H) and quantitation (I) of CD44 and CD62L expression on WT (black) and KO (blue) Rpl18+ cells. (**J**) MFI of CD44 expression of Rpl18 tet+ cells. (**K**) Flow plots of WT (black) and KO (blue) PD-1 and Tim-3 expression. (**L**) Quantification of PD-1 vs. Tim-3 expression on Rpl18+ cells. (**M**) Frequency of TCF-1^+^ in the Rpl18^+^ cells. (**N**) Histogram of TCF-1 expression in KO and WT Rpl18^+^ cells. (**O-P**) Flow plots (O) and quantification of Ki67 expression in WT (black) and KO (blue) CD8 T cells. (**Q**) TDLN cells were restimulated for 6hrs with plate-coated anti-CD3 (2 μg/ml); data are gated on CD44+ cells. Quantified cytokine expression of KO and WT activated cells within the TDLN. (**R**) Flow plots of both WT (black) and KO (blue) of TNF and IFNγ expression. Students T-test statistical analysis applied for statistical comparison (p: *<0.05, **<0.01, ***<0.001, ****<0.0001).

### Impact of TrIP deficiency on anti-tumor T cell transcriptome and TCR diversity

We next wanted to investigate the T cell-intrinsic molecular effects of CD8-specific TrIP KO during an anti-tumor response. To do this, we sorted antigen-specific (Rpl18*) cells from both tumor and TDLN at day 10 after MC38 implantation and performed both bulk RNA sequencing and TCR sequencing. Unbiased clustering of the total transcriptomes revealed separation by both tissue source and genotype (**figure S1**). Overall, there were more differentially expressed genes that cleared the threshold for significance in the WT vs. TrIP KO tumor-derived samples, compared with the draining lymph node (**figure 6A, figure S1**). Some specific genes of interest, related to the more effector-like phenotype of TrIP KO T cells discussed above, are shown in the heatmap in **figure 6B**. We next performed gene set enrichment on the differentially expressed genes, comparing WT and KO within either the tumor or draining lymph node samples. Not surprisingly, this yielded fewer significantly different hallmark gene sets than the *in vitro* stimulation discussed in **figure 1**. Nonetheless, we did find several enriched gene sets within the tumor (**figure 6C**). These included positive upregulation of mTORC1 signaling (which is downstream of PI3K) in KO cells, as well as significant enrichment of several metabolic pathways. Interesetingly several of the latter involved postitive enrichment of pathways generally involved with lipid metabolism (xenobiotic, fatty acid and bile acid metabolic pathways) in KO T cells, as well as negative enrichment in the KO cells for oxidative phosphorylation. Individiual genes that contributed to these differences included *Idh1*, *Fasn*, *Cd36* and *Fads1*. Overall, this finding is consistent with the concomitant upregulation of MTORC1 signaling in the same cell population. Within the tumor draining lymph node, very few pathways were eniched (**figure 6D**), including positive enrichment in the KO cells for KRAS siganling. Interestingly, there was a modest negative enrichement for fatty acid metabolism in the KO cells, although this was not statistically significant.

**Figure 6:**
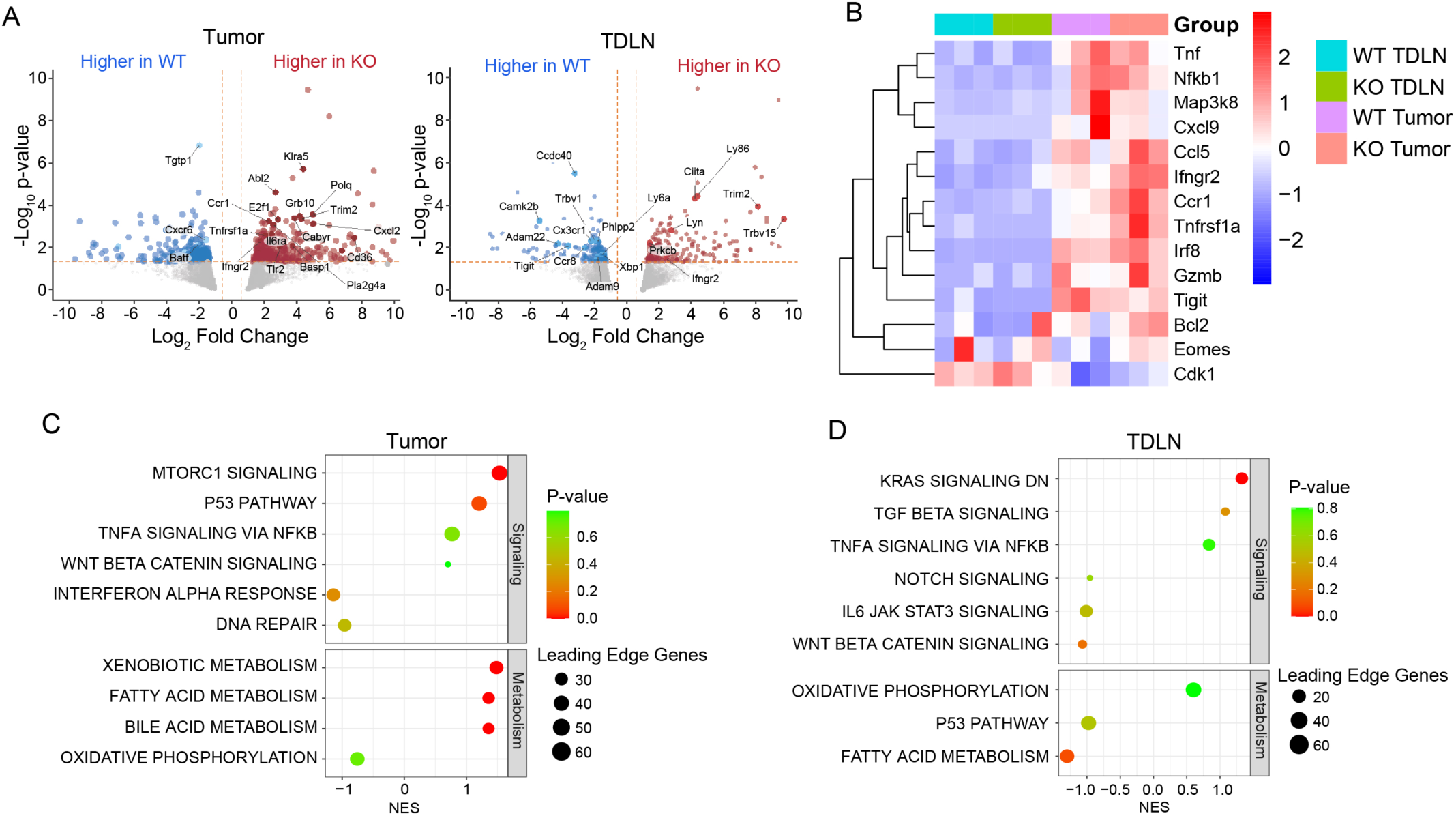
RNAseq analysis of TrIP KO vs. WT tumor neoantigen-specific CD8 T cells. WT (E8i-Cre only) or TrIP KO (E8i-Cre x *Pik3ip1^fl/fl^*) mice were implanted with MC38 tumors. On day 10, single cell suspensions were made from tumors and tumor-draining lymph nodes. Rpl18*-specific CD8 T cells were sorted by flow cytometry and processed for RNA sequencing. (**A**) Volcano plots of WT vs. TrIP KO cells from tumor (left) and tumor-draining lymph node (TDLN; right), with selected genes highlighted. (**B**) Heat map of relative expression of selected genes in individual samples. (**C-D**) Gene set enrichment results of selected pathways in T cells from tumor (C) and draining lymph node (D). A positive enrichment score indicates enrichment in the KO cells.

We next performed TCR repertoire analysis of TCR sequencing data obtained from the same samples analyzed for bulk RNA sequencing, using the TRUST4 package. Through this analysis we identified 2000-3000 unique CDR3’s per sample, with roughly 10-14% fully matched TCR’s (paired TCRα/β) (**figure 7A**). After normalizing the TCR frequencies, the WT samples were significantly less clonally diverse in both the tumor and draining lymph node samples across both TCRα and TCRβ, compared with the TrIP KO T cells (**figure B-C, figure S2**). For example, the WT α and β CDR3’s showed single clones making up over 60% of the normalized TCR frequency in these samples, compared to the highest frequency TCR in the KO representing just ∼20% (**figure 7C-D**). These findings suggest that a more diverse repertoire of T cells/TCRs may be able to clear the signaling threshold for full activation in the absence of TrIP.

**Figure 7:**
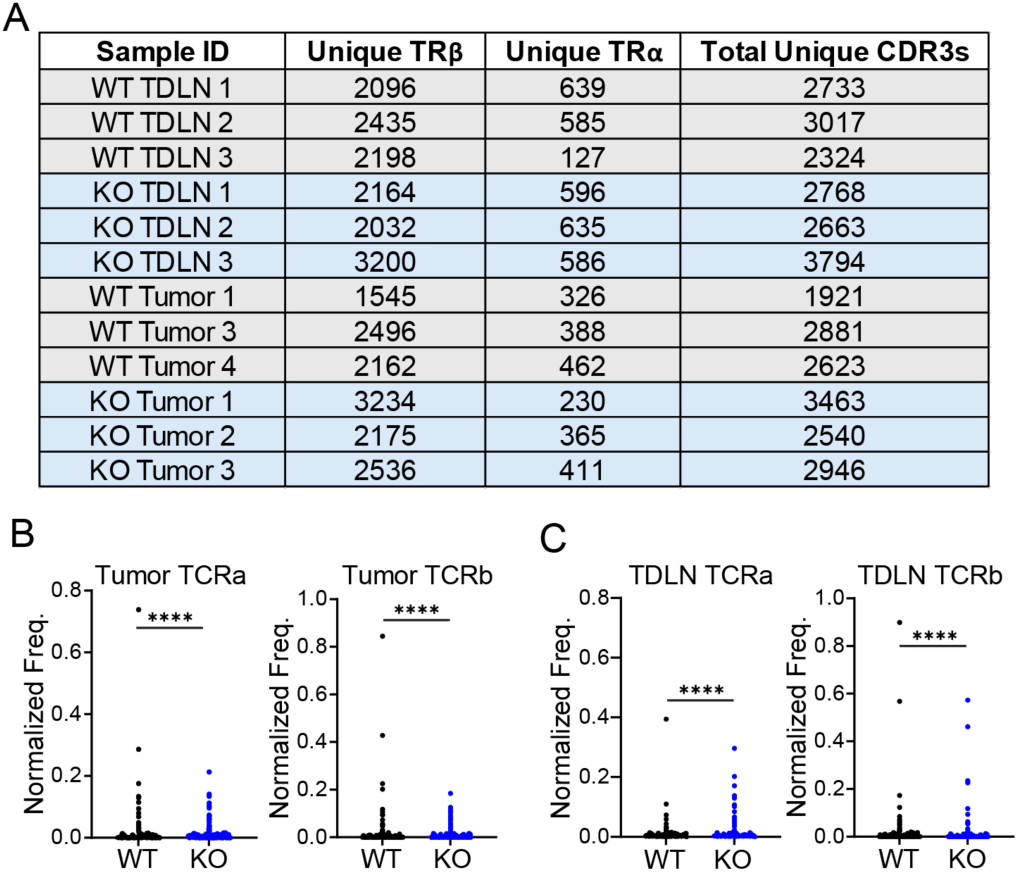
More-diverse pool of TCRs among responding anti-tumor CD8 T cells in TrIP KO mice. Samples analyzed in Fig. 6 were processed in parallel for TCRseq. TRUST4 analysis was used to determine the number of unique TCRa and TCRb transcripts in WT vs. TrIP KO Rpl18*-reactive CD8 T cells. (**A**) Tabulation of the number of unique TCRa and TCRb sequences recovered from each sample, as well as the number of unique CDR3s. (**B-C**) Normalized TCR frequencies in tumors (**A**) and tumor-draining lymph node (**B**).

## Discussion

In this study we demonstrate that TrIP expression on naïve CD8 T cells restricts early TCR activation following antigen recognition. Thus, using CD8-specific TrIP KO mice, we show that in the absence of TrIP, CD8 T cells acquire a more inflammatory transcriptional profile. Furthermore, we show that these transcriptional changes in the TrIP KO CD8 T cells are accompanied by an improvement in the immune response to syngeneic tumors, leading to a decrease in tumor burden. In addition, the absence of TrIP in CD8 T cells resulted in a broader clonal response, presumably to tumor antigens, ultimately leading to an increase in the proportion of tumor-specific CD8 T cells cells within the TrIP KO TIL.

It was not entirely unexpected that knockout of a negative regulator of PI3K would promote a more inflammatory response. However, it is still not understood how TrIP regulates CD8 T cells more broadly. Thus we initially focused on understanding in detail the transcriptional changes that occur in the absence of the early regulation of PI3K conferred by TrIP in the resting state. Indeed, RNAseq analysis of *in vitro*-activated T cells revealed that the absence of TrIP expression on naïve cells led to significant transcriptional changes in CD8 T cells, across all three treatment groups (unstimulated, stimulated +/- PI3Ki). Our finding of global changes in gene expression, even in the absence of TCR stimulation, was surprising, considering that we have not observed obvious changes in the gross phenotype of these cells by flow cytometry. Nonetheless, it is important to note that even the cells in the unstimulated condition were cultured in the presence of cytokines (IL-2 + IL-12) for the entire 24 hours prior to sequencing. IL-2 receptor signaling is known to activate the PI3K pathway, which may explain some of the differences seen under those conditions [23]. In any case, the largest differences in gene expression in TrIP KO CD8 T cells were seen after TCR stimulation, promoting more inflammatory gene expression. Building on previous germline TrIP knockout studies, these data led us to explore how the knockout of TrIP on CD8 T cells was sufficient to improve resistance to implanted syngeneic tumors.

To further characterize how the absence of TrIP imparts this resistance to tumor challenges, we investigated the tumor immune infiltrate of WT (E8i^cre^) and TrIP KO (*Pik3ip1^fl/fl^ E8i^cre^*) T cells, using a floxed mouse model that we previously described [14]. This distinguishes our study from a previous report, wherein the authors examined the effects of germline TrIP deficiency [15]. Thus, when looking at day 14 after tumor inoculation, we found smaller tumor burden and higher T cell frequencies in the TrIP^CD8^ knockout animals. This effect was associated with a selective expansion of the CD8 compartment in the tumors, demonstrating a cell-intrinsic effect of TrIP deficiency in the CD8 T cell compartment. Among the TIL CD8 T cells lacking TrIP there was a significantly lower frequency of TCF-1^+^ cells, suggesting that the KO cells were more terminally differentiated, at least at this timepoint. Previous reports have shown TCF-1 expressing cells in the TIL to be more beneficial to an effective tumor-immune response, which is not consistent with the finding in TrIP KO mice [24]. These conflicting observations could be explained by the increased number of CD8 T cells within the KO TIL. Thus, while we may have observed a reduction in the proportion of TCF-1^+^ cells (∼40%^+^ in WT vs. ∼20%^+^ in KO), the total numbers of TCF-1^+^ CD8 T cells within WT and KO TIL were actually closer. Additionally, we found a modest, but statistically significant, increase in TNF production upon restimulation of TrIP KO vs. WT T cells. Furthermore, differences in inflammatory cytokine production may also contribute to the reduction in tumor burden we observed in CD8-specific TrIP KO animals.

While the above-discussed phenotype of TrIP KO CD8 TIL T cells suggests a mechanism for the reduced tumor burden, it was at first puzzling that we did not observe TrIP protein expression on WT CD8 T cells in the tumor (data not shown). However, as we recently reported, the highest TrIP expression is found on resting/naïve T cells, and TCR stimulation leads to active cleavage of the protein from the cell surface [13]. Although TrIP expression eventually recovers, tumor-infiltrating T cells are enriched for tumor antigen specificity, leading to chronic T cell stimulation. Thus, we hypothesized that the increased CD8 T cell presence we reported in the day 14 TIL may be due to differences in initial T cell priming. To assess early responses in antigen-specific cells, we employed tetramers against a dominant neoantigen expressed by MC38, derived from ribosomal protein Rpl18 [16]. Rpl18-specific cells comprised ∼0.5-2% of the CD8 T cells in the tumor-draining lymph nodes (TDLN), but there were no detectable differences in frequency between the TrIP KO and WT at the D14 timepoint. Additionally, the WT and KO Rpl18^+^ cells largely shared the same gross phenotype in the TDLN, as they expressed similar levels of activation (CD44/CD62L) and exhaustion markers (PD-1/Tim-3). In fact, we did not observe a difference in TCF-1 expression in WT vs. KO CD8 T cells within the TDLN. We thus turned our focus towards earlier timepoints.

At day 10, tumors in WT vs. CD8-specific TrIP KO mice were of similar size; this was expected, as day 10 is prior to the bifurcation in the tumor growth curves (**Fig. 2**). In addition, TrIP KO and WT TIL T cells were found at similar frequencies in the tumors, with similar proportions of CD4 vs. CD8 T cells. However, the KO CD8 T cells appeared less exhausted phenotypically, compared with WT CD8 T cells. Lower levels of exhaustion may support the survival/persistence or proliferation of these cells as the tumors progress, which could help to explain the increased T cell infiltration and function we observed at later time points. Although we did not observe major differences in proliferation based on Ki67 staining, it is possible that a modest such difference could explain the enhanced T cell infiltration at later time points, since differences in proliferation can compound over time. Further in-depth longitudinal analyses will be necessary to answer this question more definitively.

Investigation of the response in the tumor-draining lymph nodes at day 10 revealed a small but statistically significant increase in CD8 T cell frequency in TrIP KO animals. Interestingly, this did not appear to be the result of a more robust response to the mutated Rpl18 antigen. We thus hypothesized that the absence of TrIP on naïve T cells may allow for responses to a broader range of tumor antigens that are otherwise too weak to trigger full T cell activation. Further analysis of the Rpl18 tetramer-specific cells within the TDLN revealed a phenotype reminiscent of central memory T cells, with a greater proportion of TrIP KO cells expressing both CD62L and CD44. Although there was a lower frequency of TrIP KO CD8 T cells that were CD44 single-positive, these cells had significantly higher levels of CD44 on a per-cell basis. Differences in the level of CD44 are biologically relevant, as CD44 not only identifies previously activated T cells, but can also promote signal transduction through Src family kinases and PI3K [25]. Persistent expression of CD62L on the KO T cells may help to explain the increased proportion of CD8 T cells in the TrIP KO TDLNs at day 10, as CD62L promotes T cell retention in the lymph node. We also observed higher in TCF-1 expression in the TrIP KO cells in the TDLN at day 10. Tumor-specific CD8 T cells within the TDLN that are able to maintain TCF-1 have been reported to serve as a ‘reservoir’ for replenishing the tumor-responding effector T cells, enabling prolonged/persistent anti-tumor response [26].

Based on the day 10 phenotypic data, we performed bulk RNA-seq and TCR-seq on Rpl18-specific CD8 T cells from the TDLN and TIL. Not surprisingly, there were fewer significant differences in gene expression, compared with the *in vitro* RNA-seq dataset discussed above. These differences may have resulted in part from the use of P14 TCR Tg T cells in the *in vitro* experiment, compared with the more complex native T cell repertoire analyzed *in vivo*. In addition, they could be due to the different *in vivo* vs. *in vitro* stimulation conditions, resulting in different nutrient/cytokine availability. Furthermore, our TCR sequencing analysis revealed that the absence of TrIP leads to a more diverse clonal response in both the TDLN and the TIL of the TrIP KO CD8 T cells. We propose that this broader response is in part responsible for the increased frequency of CD8 TIL we observed at the later (day 14) timepoint. In addition, thbe more diverse response at day 10 TIL may contribute to the lower overall exhaustion marker expression (PD-1^+^Tim3^+^) by TrIP KO CD8 T cells. This is in line with previous reports that broader repertoires may delay the progression towards exhaustion, at least in chronic viral infection models [24].

While the majority of immunotherapies in recent times have focused on targeting immune checkpoint molecules whose expression is highly induced following activation, TrIP targeting offers the potential to target and trigger separate populations of cells than those being targeted by the major immunotherapies. Along these lines, targeting of other members of the PI3K pathway, e.g. Akt and mTOR, has also been explored in settings of immunotherapy. Although partially inhibiting these kinases enhanced the acquisition of a memory T cell phenotype, this approach also limited T cell proliferation [27]. Based on a recent study of the PI3K regulator BCAP in regulating effector versus memory differentiation of CD8 T cells [28], among other studies, one way to improve CAR T cell therapy may be the identification of the right strategy to optimally “tune” PI3K signaling [28–30]. Thus, therapeutic targeting of TrIP in combination with established immunotherapy modalities may represent a novel approach for improving patient outcomes.

## Contributors

BM planned and carried out the experiments and performed analysis of data; he also generated the figures, wrote the draft of the manuscript, and obtained funding. LK conceived of the project, obtained funding, assisted with data analysis and interpretation and edited the manuscript.

## Data availability statement

All data relevant to the study are included in the article, uploaded as supplementary information or available at the Gene Expression Omnibus (GEO; https://www.ncbi.nlm.nih.gov/geo/), accession numbers GSE302184, GSE302185 and GSE302196.

## Acknowledgements

We would like to thank Grace Bowman and Edgar Cardona for their help with mouse genotyping. We thank Dominic Golec for advice on GSEA of FOXO1-regulated genes, and Alok Joglekar and Emily Landy for advice on TCRseq analysis. We also gratefully acknowledge the assistance of the Univeristy of Pittsburgh Unified Flow Cytometry Core and the Health Sciences Library. RNA sequencing library generation and sequencing were performed in the Health Sciences Sequencing Core at UPMC Children’s Hospital of Pittsburgh, Rangos Research Center. Services and instruments used in this project were supported, in part, by the University of Pittsburgh, the Office of the Senior Vice Chancellor for Health Sciences, the Department of Pediatrics, the Institute for Precision Medicine, and the Richard K Mellon Foundation for Pediatric Research (RRID:SCR_023116).

