## Supplemental Figures for "Pik3ip1/TrIP Regulation of PI3K Restricts CD8 T Cell Anti-Tumor Immunity"

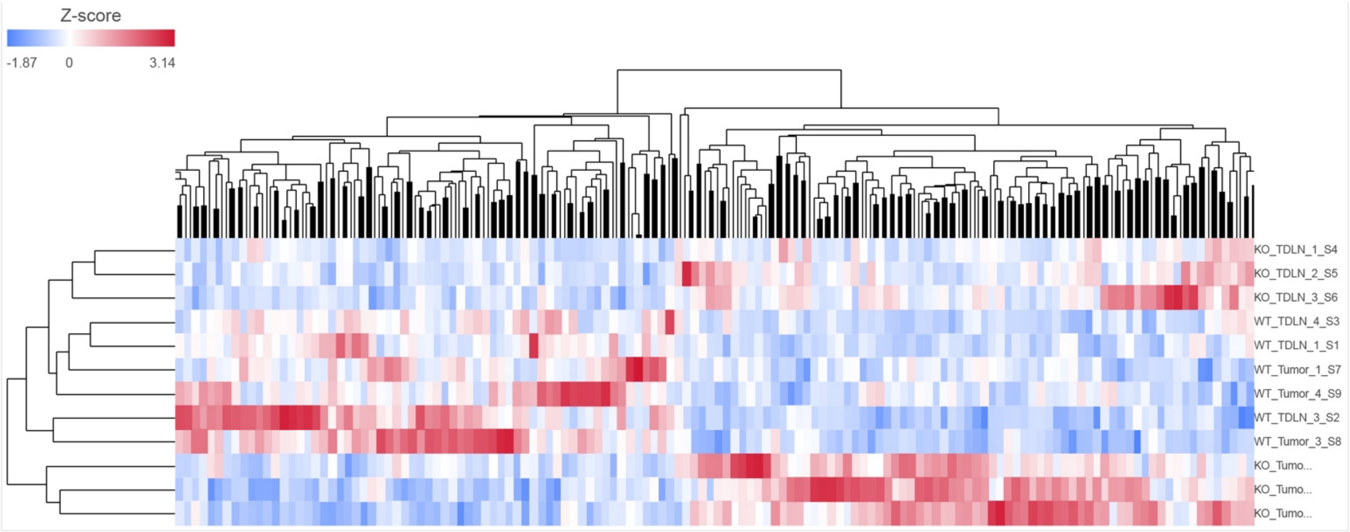


**Fig. S1: Clustering of transcripts from Rpl18*-specific CD8 T cells isolated from tumor and tumor-draining lymph node (TDLN).** DEseq2 Diff analysis was applied to all samples, comparing TrIP KO and WT (irrespective of tissue). Resulting genes were filtered p<.05 and fold change excluded 0.5-2 = ~300 DE genes globally.

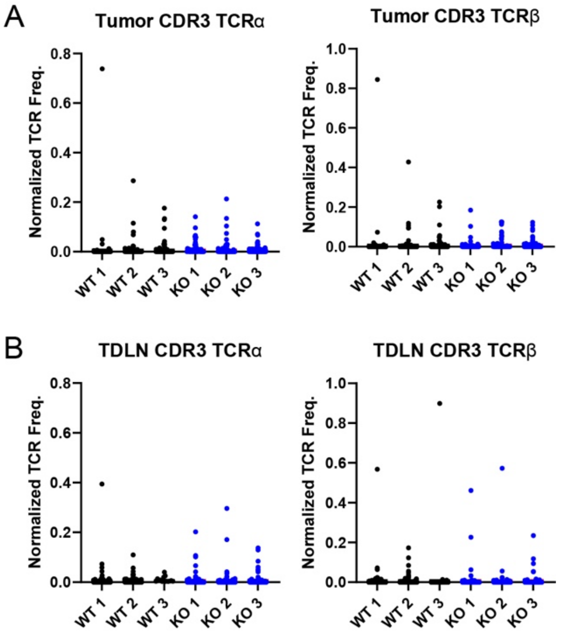


**Fig. S2: Frequencies of TCRα and TCRβ CDR3s from individual samples.** Normalized frequencies obtained from Rpl18*-reactive T cells sorted from individual tumors (**A**) and tumor-draining lymph nodes (**B**). See also Figure 7.
